# Bioactivity-guided isolation of rosmarinic acid as a principle bioactive compound from the butanol extract of *Isodon rugosus* against pea aphid, *Acyrthosiphon pisum*

**DOI:** 10.1101/591271

**Authors:** Saira Khan, Clauvis Nji Tizi Taning, Elias Bonneure, Sven Mangelinckx, Guy Smagghe, Raza Ahmad, Nighat Fatima, Muhammad Asif, Mohammad Maroof Shah

## Abstract

Aphids are agricultural pest insects that transmit viruses and cause feeding damage on a global scale. Current pest control involving the excessive use of synthetic insecticides over decades has led to multiple forms of aphid resistance to most classes of insecticides. In nature, plants produce secondary metabolites during their interaction with insects and these metabolites can act as toxicants, antifeedants, anti-oviposition agents and deterrents towards the insects. In a previous study, we demonstrated that the butanol fraction from a crude methanolic extract of an important plant species, *Isodon rugosus* showed strong insecticidal activity against the pea aphid, *Acyrthosiphon pisum*. It was however not known as which compound was responsible for such activity. To further explore this finding, current study aimed to exploit a bioactivity-guided strategy to isolate and identify the active compound in the butanol fraction of *I. rugosus*. As such, reversed-phase flash chromatography, acidic extraction and different spectroscopic techniques were used to isolate and identify the new compound, rosmarinic acid as the bioactive compound in *I. rugosus*. Insecticidal activity of rosmarinic acid was carried out using standard protocols on *A. pisum*. The data was analyzed using qualitative and quantitative statistical approaches. Considering that a very low concentration of this compound (LC_90_ = 5.4 ppm) causes significant mortality in *A. pisum* within 24 h, rosmarinic acid could be exploited as a potent insecticide against this important pest insect. Furthermore, *I. rugosus* is already used for medicinal purposes and rosmarinic acid is known to reduce genotoxic effects induced by chemicals, hence it is expected to be safer compared to the current conventional pesticides. While this study highlights the potential of *I. rugosus* as a possible biopesticide source against *A. pisum*, it also provides the basis for further exploration and development of formulations for effective field application.

## Introduction

Aphids are among the most important agricultural pest insects of many crops worldwide. They feed exclusively on plant phloem sap by inserting their needle-shaped mouthparts into sieve elements, usually resulting to the stunting, discoloration and deformation of plants, while the growth of sooty molds on honeydew produced by these insects reduces the economic value of crops [1, 2]. Moreover, aphids are also vectors of many important plant viruses [3-5]. The pea aphid, *Acyrthosiphon pisum* (Hemiptera: Aphididae), adversely affects economically important legume crops worldwide. It is oligophagous, comprising of a number of biotypes or races living on a number of legume hosts (red clover, pea and broad bean and alfalfa races) [6-9]. Current aphid control strategies predominantly rely on the use of insecticides such as carbamates, organophosphates, pyrethroids, neonicotinoids and pymetrozine [10]. However, the frequent use of these insecticides over the decades has led to multiple forms of aphid resistance to most classes of insecticides, making it very difficult to control this insect pest [11].

The use of botanical pesticides could present a safe alternative compared to the use of broad spectrum chemical insecticides in crop protection [12, 13]. In nature, plants produce secondary metabolites during their interaction with insects and these metabolites can act as toxicants, antifeedants, anti-oviposition agents and deterrents towards the insects [14-16]. Because of such wide insecticidal properties, the study of secondary metabolites and the development of new potent formulations based on them has become increasingly important. Screening of plant extracts followed by bioactivity-guided fractionation, isolation and identification of active principles is considered to be one of the most successful strategies for the discovery of bioactive natural products against insect pests [17].

*Isodon rugosus* (Wall. ex Benth.) Codd (syn. *Plectranthus rugosus* Wall. ex. Benth.) is an aromatic branched shrub, belonging to the family Lamiaceae. The plant is used in Pakistani traditional medicine for many diseases as an antiseptic, hypoglycemic, antidiarrheal and as bronchodilator [18, 19]. Among many other traditional medicinal uses, the plant extracts and different solvent fractions are known to be effective as antifungal, antibacterial, phytotoxic and antioxidant agents and are able to show lipoxygenase inhibitory activities [20-23]. Based on phytochemical studies, this plant is known to contain steroids, flavonoids, terpenoids, saponins, tannins, cardiac glycosides, coumarins, reducing sugars and β-cyanin. Diterpenoids (rugosinin, effusanin-A, effusanin-B, effusanin-E, lasiokaurin and oridonin) and triterpenoids (plectranthoic acid A and B, acetyl plectranthoic acid and plectranthadiol) have also been successfully isolated from this plant [24-26]. However, despite several studies on the bioactivity of *I. rugosus* where most efforts were focused towards human health, none of these have isolated and assessed the insecticidal activity of compounds from this plant. In a previous study, we evaluated the aphicidal properties of the hexane, dichloromethane, butanol and ethyl acetate fractions of a crude methanolic extract from *I. rugosus*, and confirmed that the butanol fraction showed the best activity against the pea aphid, *A. pisum* [27]. To further explore this finding, a bioactivity-guided strategy against *A. pisum* was used to isolate and identify the active compound in the butanol fraction of *I. rugosus*.

## Materials and methods

### Insects

A continuous colony of *A. pisum* was maintained on faba bean plants (*Vicia faba*) in the Laboratory of Agrozoology at Ghent University, Belgium at 23–25 °C and 65±5% relative humidity (RH) under a 16:8 h light: dark photoperiod [28]. All the bioassays were performed under these conditions. Newly born nymphs (< 24 h old) of *A. pisum* were used for all the bioassays. By gentle probing of the aphids with a brush and also by observing post-mortem color change of the body, mortality was assessed after 24 h of treatment.

### Plant collection and extraction

The aerial parts of *I. rugosus* were collected from lower Northern areas of Pakistan in the month of October, 2012. The plant material was shade-dried for up to 3 months and ground to powder using an electric grinder. 1 kg of the dried powder was soaked in a glass jar containing 3 L of methanol at room temperature. After two days, the solvent layer was filtered with Whatman filter paper No. 1 and this process was repeated three times. The resulting filtrate was concentrated by using a rotary evaporator at 35 °C and the obtained crude methanolic extract was stored at 4 °C [29, 27]. For fractionation, 90 g dried crude methanolic extract was mixed with five parts of water and then extracted successively by n-hexane (4 × 150 mL), dichloromethane (4 × 150 mL), ethyl acetate (4 ×150 mL) and n-butanol (4 × 150 mL) as described by Khan et al. [27]. All the fractions were concentrated using a rotary evaporator under reduced pressure at 40 °C. The resulting extracts were stored in a refrigerator at 4 °C until further use.

### Isolation of the bioactive principle

Based on bioassays conducted by Khan et al, [27] the butanol extract presented the best biological activity against *A. pisum* and was hence selected in this study for further bioactivity-guided fractionation and identification of the active principle. The butanol extract (500 mg) was eluted with a Reveleris automated flash chromatography instrument on a 12 g C18 pre-packed column (GRACE, Columbia, MD, US) starting with 100% water. The gradient was ramped to 100% methanol over 60 column volumes (CV) and after collection of 95 fractions, the solid phase was flushed with 5 CV acetonitrile. The flow rate was set to 30 mL/min (Table 1). Based on the UV spectral data, the 95 fractions were combined into a total of 14 subfractions. These combined fractions were evaporated under reduced pressure at 45 °C and finally under a high vacuum resulting in 14 subfractions (1A-14A) (Table 2). The 14 subfractions were evaluated for their bioactivity against *A. pisum*, of which fraction 3A was selected on the basis of maximum bioactivity for further fractionation through preparative liquid chromatography (prep-LC). A 10% solution of fraction 3A was prepared in methanol. Two solvents were used, water (solvent A) and acetonitrile (solvent B). A gradient was set starting with 100% solvent A from 0 to 100 min. From 100 min to 110 min, solvent B went from 18% to 100% and stayed at 100% until 128 min, and then to 0% at 128.10 min and stayed at 0% until min 132.10 min. After concentration under reduced pressure with a rotary evaporator and finally under a high vacuum, three fractions, 3A-1, 3A-2 and 3A-3 were obtained. Fraction 3A-3 was selected for active compound identification (NMR and LC-MS) on the basis of the bioactivity against *A. pisum*. This compound was obtained in pure form by doing a second flash chromatographic separation of 5 g of butanol extract and by using the run conditions as mentioned in Table 3. From the second flash chromatography, a total of 354 fractions were collected which were combined into six fractions, 1B, 2B, 3B, 4B, 5B and 6B on the basis of UV spectra and were further analyzed for their bioactivity after concentration with a rotary evaporator under reduced pressure and high vacuum (Table 4). Fraction 1B was selected for further purification on the basis of best bioactivity. On the basis of knowledge regarding the acidic compound present in sub fraction 3A-3 (from ^1^H NMR and HPLC-MS analysis), an extraction under acidic conditions was done to isolate the active compound from sub fraction 1B. For this purpose, 200 mg of fraction 1B was dissolved in 10 mL of distilled water and acidified with 4 drops of hydrochloric acid (12 M). Following extraction with ethyl acetate (four times 5 mL), two phases, ethyl acetate and aqueous, were obtained. Both the ethyl acetate and the aqueous phase were concentrated. The ethyl acetate phase fraction was more bioactive with lower LC values. Last traces of ethyl acetate were removed azeotropically with toluene and evaporation under high vacuum of the residues resulted in 60 mg from the ethyl acetate phase and 60 mg from the aqueous phase. The purified active principle was identified through different spectroscopic techniques.

**Table 1.** First reversed-phase flash chromatography conditions of butanol fraction (500 mg) from *Isodon rugosus*

| **Run Conditions** |  |
| --- | --- |
| **Cartridge** | Reveleris 12 g C18 40 µm |
| **Solvent A** | Water |
| **Solvent B** | Methanol |
| **Solvent C** | Acetonitrile |
| **Flow rate** | 30 mL/min |
| **Injection type** | Dry sample |
| **ELSD Carrier** | Isopropanol |
| **Per vial volume** | 25 mL |
| **UV1 Wavelength** | 220 nm |
| **UV2 Wavelength** | 254 nm |

**Table 2.** Subfractions (1A-14A) collected from the first reversed-phase flash chromatography of butanol extract (500 mg)

| Fractions | Weight (mg) |
| --- | --- |
| 1A | 52 |
| 2A | 11 |
| 3A | 46 |
| 4A | 7 |
| 5A | 20 |
| 6A | 15 |
| 7A | 54 |
| 8A | 58 |
| 9A | 14 |
| 10A | 49 |
| 11A | 18 |
| 12A | 15 |
| 13A | 21 |
| 14A | 18 |

**Table 3.** Second reversed-phase flash chromatography conditions of butanol fraction (5 g) of *Isodon rugosus*

| Run Conditions |  |
| --- | --- |
| Cartridge | Reveleris 120 g C18 40<br>µm |
| Solvent A | Water |
| Solvent B | Methanol |
| Flow rate | 85 mL/min |
| Injection type | Dry sample |
| ELSD Carrier | Isopropanol |
| Per vial volume | 25 mL |
| UV1 Wavelength | 220 nm |
| UV2 Wavelength | 254 nm |

**Table 4.** Subfractions (1B-6B) from the second reversed-phase flash chromatography of butanol extract (5 g)

| Fractions | Weight (mg) |
| --- | --- |
| 1B | 530 |
| 2B | 830 |
| 3B | 1523 |
| 4B | 195 |
| 5B | 140 |
| 6B | 128 |

### Identification of the bioactive compound

Mass spectra were recorded using a HPLC-MS instrument consisting of an Agilent (Waldbronn, Germany) model 1100 liquid chromatograph with a diode array detector coupled with a mass spectrometer with electrospray ionization geometry (Agilent MSD 1100 series). The prep-LC consisted of an Agilent 1100 Series liquid chromatograph using a Supelco Ascentis C18 column (I.D. × L 21.2 mm × 150 mm, 5 µm particle size) connected to an UV-VIS variable wavelength detector (VWD) and automatic fraction collector. Flash chromatography was performed with the Reveleris Flash System (GRACE). ^1^H and ^13^C NMR spectra were obtained on a BRUKER Advance III 400 spectrometer. All the solvents and chemicals used were of analytical grade. Optical rotation was taken with a JASCO P-2000 series polarimeter.

### Insecticidal bioactivity

For the bioassays, artificial diet test cages were prepared as described by Sadeghi et al. [30] 100 µL of liquid artificial diet was sealed between two layers of parafilm. Ten neonate aphids were placed on these layers of the parafilm and to prevent the escape of aphids, the cages were covered with a hollow plastic ring having a ventilated lid. These cages were placed in an inverted position in six aerated well plates. Five concentrations were used for each treatment against the aphids. A stock solution of 1% was prepared by adding 1 mg of each fraction in 100 µL of water. In case of the reversed-phase flash fractions, five concentrations with 50, 25, 12.5, 6.3 and 3.1 ppm and in case of the prep-LC and acidic extraction fractions, five concentrations of 5, 2.5, 1.3, 0.7 and 0.3 ppm were prepared by diluting the stock solution with the artificial diet of aphids. For each concentration, a final volume of 300 µL was made to carry out three replications of each treatment (100 µL for each replication). Pure isolated and identified active compound was analyzed in eight different concentrations including 50, 25, 12.5, 6.3, 3.1, 1.6, 0.8 and 0.4 ppm by using a stock solution of 1 mg of compound in 100 µL of water. The untreated artificial diet was used as a control and three replications were used for each treatment in all the bioassays. Mortality was analyzed after 24 h of each treatment.

Additionally, the growth of the surviving aphids exposed to 0.4 ppm of the active compound for 24 h was followed for 9 days (on the same treated diet) in comparison to the untreated aphids.

### Data analysis

For statistical analysis, Probit analysis of mortality vs. concentration using POLO-Plus program version 2 was conducted and the lethal concentrations (LC_50_, LC_90_) and their corresponding 95% confidence intervals (95% CI) were estimated for each fraction. When the 95% CI’s did not overlap, LC’s were considered to be significantly different.

## Results

### Bioactivity of fractions from the butanol extract of ***I. rugosus***

Bioactivity of the fourteen fractions (1A-14A) obtained through the first reversed-phase flash chromatography of 500 mg of butanol extract of *I. rugosus* was analyzed for 24 h against *A. pisum*. Except fractions 8A, 9A, 11A, 13A and 14A, all the other fractions showed considerable toxic effects against *A. pisum*. Fraction 3A (LC_50_ = 2.1 ppm and LC_90_ = 29.5 ppm) had the highest activity as compared to all other fractions, followed by fraction 5A (LC_50_ = 3.3 ppm and LC_90_ = 50 ppm). Fraction, 1A (LC_50_ = 5.5 ppm and LC_90_ = 66 ppm), 2A (LC_50_ = 8.9 ppm and LC_90_ = 81 ppm), 4A (LC_50_ = 6.8 ppm and LC_90_ = 112 ppm) and 6A (LC_50_ = 17.8 ppm and LC_90_ = 187 ppm) gave considerable mortality. Fraction 7A (LC_50_ = 74 ppm and LC_90_ = 267 ppm) showed lower mortality. Moderate toxicity was observed with fraction 10A (LC_50_ = 36 ppm and LC_90_ = 53 ppm) and 12A (LC_50_ = 51 ppm and LC_90_ = 109 ppm) (Table 5).

**Table 5.** Toxicity of subfractions of the butanol fraction from first reversed-phase flash chromatography against newborn (< 24 h old) *Acyrthosiphon pisum nymphs* following 24 h exposure to artificial diet containing different concentrations of subfractions

| Fractions | LC<sub>50</sub> (95% CI) ppm | Ratio | LC<sub>90</sub> (95% CI) ppm | Ratio | Slope ± SE | Chi-Square | HF |
| --- | --- | --- | --- | --- | --- | --- | --- |
| 1A | 5.5 (3-8) a | 2.6 | 66 (37-211) a | 2.2 | 1.1 ± 0.3 | 7.1 | 0.5 |
| 2A | 8.9 (6.1-12) a | 4.2 | 81 (47-231) a | 2.7 | 1.3 ± 0.3 | 5.6 | 0.4 |
| 3A | 2.1 (0.6-3.8) a | 1.0 | 30 (18-85) a | 1.0 | 1.1 ± 0.3 | 7.5 | 0.6 |
| 4A | 6.8 (3.8-10) a | 3.2 | 112.2 (54-561) a | 3.8 | 1.1 ± 0.3 | 4.6 | 0.4 |
| 5A | 3.3 (1.3-5.4) a | 1.6 | 50 (28-176) a | 1.7 | 1.1 ± 0.3 | 10.1 | 0.8 |
| 6A | 18 (13-27) b | 8.5 | 187 (90-808) a | 6.3 | 1.3 ± 0.3 | 3.8 | 0.3 |
| 7A | 74 (52-169) c | 35.3 | 267 (131-1651) a | 9.1 | 2.3 ± 0.6 | 8.1 | 0.6 |
| 8A | - | - | - | - | 1.7 ± 0.7 | 7.0 | 0.5 |
| 9A | - | - | - | - | 2.0 ± 1.3 | 4.7 | 0.4 |
| 10A | 36 (33-40) d | 17.2 | 52.5 (46-64) a | 1.8 | 8.0 ± 1.4 | 2.8 | 0.2 |
| 11A | - | - | - | - | 1.6 ± 0.6 | 8.5 | 0.7 |
| 12A | 51 (43-71) c | 24.5 | 109 (77-241) a | 3.7 | 3.9 ± 1.0 | 2.2 | 0.2 |
| 13A | - | - | - | - | 1.5 ± 1.2 | 6.6 | 0.5 |
| 14A | - | - | - | - | 2 ± 1.3 | 4.7 | 0.4 |

Data is presented as 50% (LC<sub>50</sub>) and 90% (LC<sub>90</sub>) lethal concentration values (both in ppm) together with their respective 95% confidence interval (95% CI), the slope ± SE of the toxicity vs concentration curve, and the Chi-Square and heterogeneity factor HF as accuracy of data fitting to probit analysis in POLO-PlusV2. Different letters in the same column indicate significant differences due to non-overlapping of 95% CI. Ratio, LCx, fraction/LCx, 3A

### Bioactivity of subfractions from fraction 3A collected through prep-LC

The three collected subfractions (3A-1, 3A-2 and 3A-3) of 3A were analyzed against *A. pisum* for 24 h. Fraction 3A-1 and fraction 3A-2 gave negligible toxic effects (no LC_50_ and LC_90_). Fraction 3A-3 was the most toxic fraction analyzed against *A. pisum* with low LC’s (LC_50_ = 1 ppm and LC_90_ = 14 ppm) (Table 6).

**Table 6.** Toxicity of subfractions of fraction 3A against newborn (< 24 h old) *Acyrthosiphon pisum* nymphs following 24 h exposure to artificial diet containing different concentrations of subfractions

| Fractions | LC<sub>50</sub> (95% CI) ppm | Ratio | LC<sub>90</sub> (95% CI) ppm | Ratio | Slope ± SE | Chi-Square | HF |
| --- | --- | --- | --- | --- | --- | --- | --- |
| 3A-1 | - | - | - | - | 2.0 ± 1.3 | 4.9 | 0.4 |
| 3A-2 | - | - | - | - | 1.5 ± 1.2 | 6.6 | 0.5 |
| 3A-3 | 1 (0.6-1.6) a | 1 | 14 (6.1-97) a | 1 | 1.1 ± 0.3 | 14.8 | 1.1 |

Data is presented as 50% (LC50) and 90% (LC90) lethal concentration values (both in ppm) together with their respective 95% confidence interval (95% CI), the slope ± SE of the toxicity vs concentration curve, and the Chi-Square and heterogeneity factor HF as accuracy of data fitting to probit analysis in POLO-PlusV2. Different letters in the same column indicate significant differences due to non-overlapping of 95% CI. Ratio, LCx, fraction/LCx, 3A-3

### Spectroscopic analysis of fraction 3A-3

Out of three subfractions of 3A (3A-1, 3A-2 and 3A-3), fraction 3A-3 was the most bioactive fraction against *A. pisum*. This fraction 3A-3 was analyzed through ^1^H NMR which confirmed that the bioactive fraction 3A-3 contained rosmarinic acid. Different gradients were used to purify the compound but during different Prep-LC runs, the chromatographic behavior, that is, peak shape and position, of this fraction was inconsistent. Therefore, the reversed-phase flash chromatography was repeated with 5 g of butanol fraction of *I. rugosus* in order to get the most bioactive compound in pure form.

### Bioactivity of fractions of butanol extract from the second reversed-phase flash chromatography

Six fractions (1B-6B) obtained through second reversed-phase flash chromatography of the butanol extract of *I. rugosus*, were analyzed against *A. pisum* for 24 h. Out of the six fractions analyzed, fraction 4B, 5B and 6B showed negligible toxicity (no LC_50_ and LC_90_). Fraction 1B was more toxic (LC_50_ = 2.5 ppm and LC_90_ = 28 ppm) and moderate toxicity was observed for fraction 2B (LC_50_ = 7.5 ppm and LC_90_ = 71 ppm). Lower toxicity was found for fraction 3B (LC_50_ = 16.3 ppm and LC_90_ = 101 ppm) (Table 7).

**Table 7.**
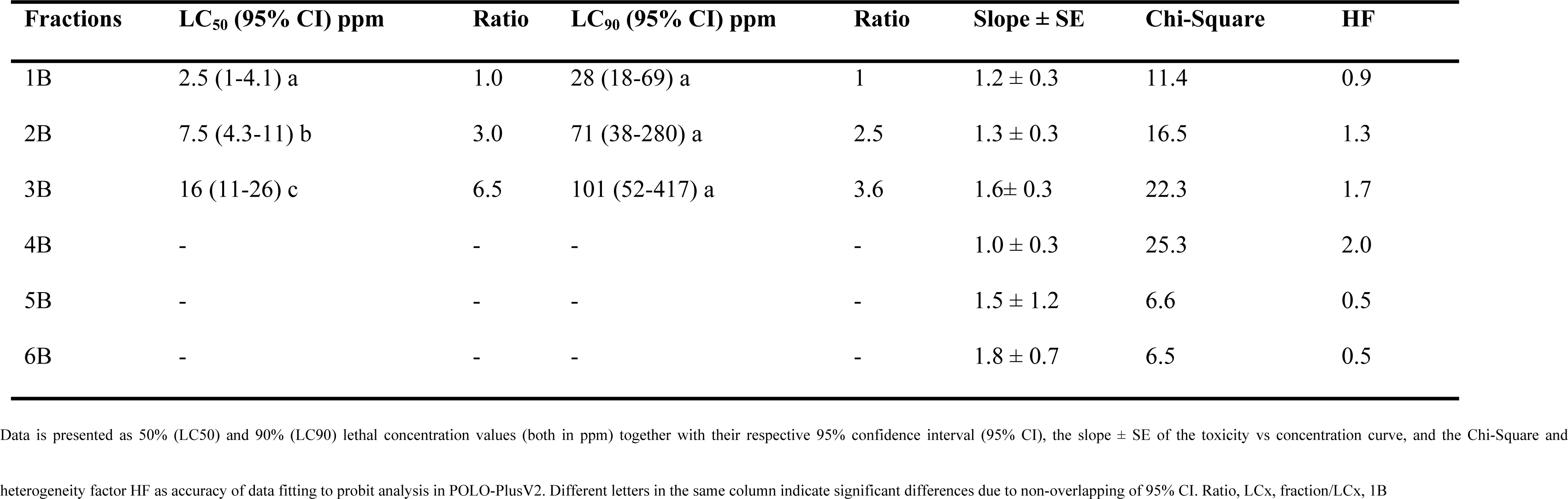
Toxicity of subfractions of the butanol fraction from second reversed-phase flash chromatography against newborn (<24 h old) *Acyrthosiphon pisum* nymphs following 24 h exposure to artificial diet containing different concentrations of subfractions

### Bioactivity of the ethyl acetate and aqueous phase of acidic extraction

Both collected phases of acidic extraction were analyzed for their insecticidal potential through bioassay against *A. pisum* for 24 h. The aqueous phase gave negligible toxic effect (no LC_50_ and LC_90_) while the ethyl acetate phase showed more toxicity (LC_50_ = 0.2 ppm and LC_90_ = 9.2 ppm) (Table 8).

**Table 8.** Toxicity of ethyl acetate and aqueous phase of acidic extraction against newborn (< 24 h old) *Acyrthosiphon pisum* nymphs following 24 h exposure to artificial diet containing different concentrations of both phases

| Fractions | LC<sub>50</sub> (95% CI) ppm | Ratio | LC<sub>90</sub> (95% CI) ppm | Ratio | Slope ± SE | Chi-Square | HF |
| --- | --- | --- | --- | --- | --- | --- | --- |
| Aqueous | - | - | - | - | 1.5 ± 1.2 | 6.6 | 0.5 |
| Ethyl acetate | 0.2 (0.04-0.5) a | 1 | 9.2 (3.9-13)a | 1 | 0.8 ± 0.3 | 4.2 | 0.3 |

Data is presented as 50% (LC50) and 90% (LC90) lethal concentration values (both in ppm) together with their respective 95% confidence interval (95% CI), the slope ± SE of the toxicity vs concentration curve, and the Chi-Square and heterogeneity factor HF as accuracy of data fitting to probit analysis in POLO-PlusV2. Different letters in the same column indicate significant differences due to non-overlapping of 95% CI. Ratio, LCx, fraction/LCx, ethyl acetate

### Identification of the most bioactive compound

Out of the two phases of acidic extraction, the ethyl acetate phase fraction was the most active. After removing ethyl acetate azeotropically, this fraction was analyzed and the active compound was identified as rosmarinic acid through HPLC-MS, optical rotation measurement and ^1^H and ^13^C NMR spectroscopy.

### HPLC-MS

Both isolated and commercial rosmarinic acid (Sigma Aldrich) had the same peak appearance in the HPLC-MS chromatograms with the same solvent gradient. Both had a pseudo-molecular ion with an *m/z* value of 359 with negative mode electrospray ionization which confirmed that it was rosmarinic acid (Fig 1).

**Fig 1.**
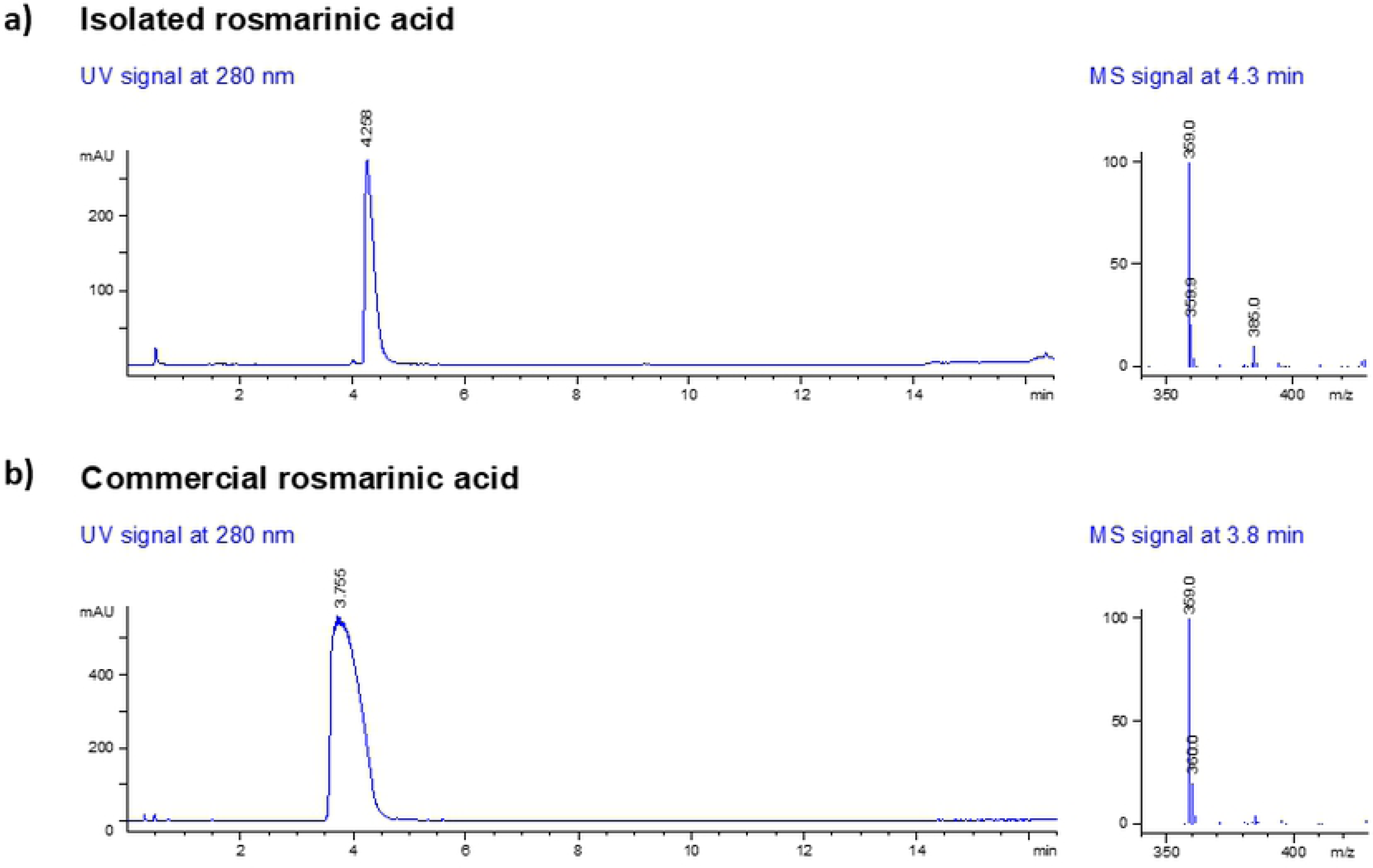
Mass spectra (negative mode electrospray ionization) of rosmarinic acid obtained via HPLC-MS with a pseudo molecular ion at *m/z* value of 359. (a) Isolated rosmarinic acid (b) Commercial rosmarinic acid

### Optical rotation and ^1^H and ^13^C NMR

Brown crystals; 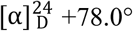 (*c* 0.233, MeOH); **^1^H NMR (400 MHz, CD_3_OD): δ**3.01 (1H, dd, *J* = 8.3, 14.3 Hz, H^7a^), 3.10 (1H, dd, *J* = 4.4, 14.3 Hz, H^7b^), 5.19 (1H, dd, *J* = 4.4, 8.3 Hz, H^8^), 6.27 (1H, d, *J* = 15.9, H^17^), 6.61 (1H, dd, *J* = 2.0, 8.0 Hz, H^6^), 6.70 (1H, d, *J* = 8.0 Hz, H^5^), 6.75 (1H, d, *J* = 2.0 Hz, H^2^), 6.78 (1H, d, *J* = 8.2 Hz, H^14^), 6.95 (1H, dd, *J* = 2.0, 8.2 Hz, H^15^), 7.04 (1H, d, *J* = 2.0 Hz, H^11^), 7.55 (1H, d, *J* = 15.9 Hz, H^16^);**^13^C NMR (100 MHz, CD_3_OD): δ** 37.9 (C^7^), 74.6 (C^8^), 114.4 (C^17^), 115.2 (C^11^), 116.3 (C^5^), 116.5 (C^14^), 117.6 (C^2^), 121.8 (C^6^), 123.2 (C^15^), 127.7 (C^10^), 129.2 (C^1^), 145.3 (C^4^), 146.2 (C^3^), 146.8 (C^12^), 147.7 (C^16^), 149.7 (C^13^), 168.4 (C^18^), 173.5 (C^9^); **ESI-MS:** *m/z* (%) 359 (M-H^+^, 100). Optical rotation and NMR data were in accordance with the literature (Fig 2) [31, 32].

**Fig 2.**
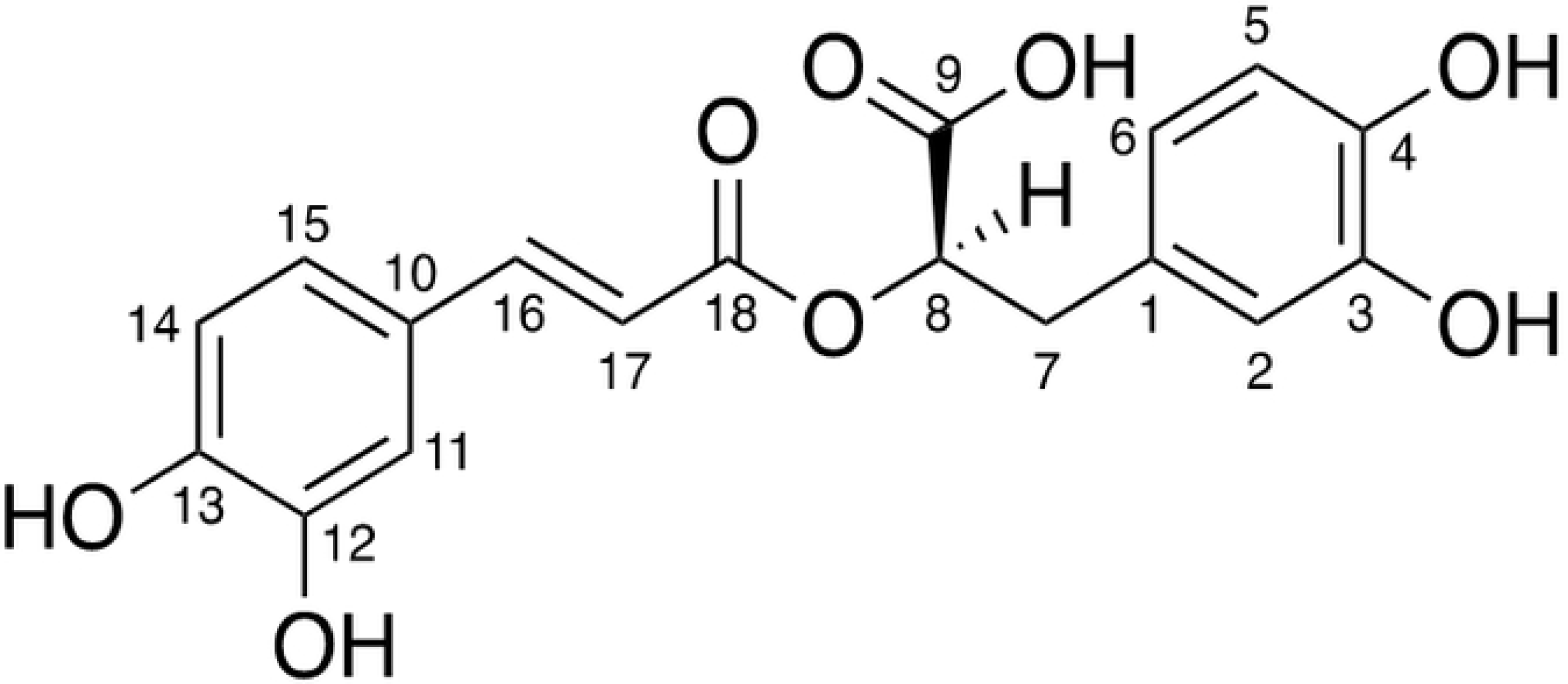
Structure of rosmarinic acid isolated from *I. rugosus*.

### Bioactivity of *I. rugosus* rosmarinic acid and commercial rosmarinic acid

Rosmarinic acid isolated from *I. rugosus* and commercial rosmarinic acid (Sigma Aldrich) were analyzed against *A. pisum* for their pesticidal activity for 24 h. Both *I. rugosus* rosmarinic acid (RA) (LC_50_ = 0.2 ppm and LC_90_ = 5.4 ppm) and commercial RA (LC_50_ = 0.2 ppm and LC_90_ = 14 ppm) gave similar toxic effects (Table 9).

**Table 9.** Toxicity of isolated rosmarinic acid (RA) and commercial rosmarinic acid (RA) against newborn (< 24 h old) *Acyrthosiphon pisum* nymphs following 24 h exposure to artificial diet containing different concentrations of isolated rosmarinic acid and commercial rosmarinic acid

| Compound | LC<sub>50</sub> (95% CI) ppm | Ratio | LC<sub>90</sub> (95% CI) ppm | Ratio | Slope ± SE | Chi-Square | HF |
| --- | --- | --- | --- | --- | --- | --- | --- |
| Commercial RA | 0.2 (0.05-0.5) a | 1 | 14 (7.4-42) a | 2.6 | 0.7 ± 0.2 | 15.5 | 0.7 |
| *I. rugosus* RA | 0.2 (0.04-0.4) a | 1 | 5.4 (3.3-12) a | 1 | 0.8 ± 0.2 | 10.5 | 0.5 |

Data is presented as 50% (LC50) and 90% (LC90) lethal concentration values (both in ppm) together with their respective 95% confidence interval (95% CI), the slope ± SE of the toxicity vs concentration curve, and the Chi-Square and heterogeneity factor HF as accuracy of data fitting to probit analysis in POLO-PlusV2. Different letters in the same column indicate significant differences due to non-overlapping of 95% CI. Ratio, LCx, compound/LCx, *Isodon rugosus* RA

### Comparison of the growth of surviving aphids exposed to rosmarinic acid-treated and untreated diet after 24 h of bioassay

After incorporating the rosmarinic acid in aphid’s diet at 0.4 ppm, its effect on *A. pisum* that survived after 24 h treatment, was analyzed every day for up to 9 days (on same treated diet). It was confirmed that rosmarinic acid had a drastic effect on their growth. Firstly, most aphids exposed to treated diet were dead while the survivors did not grow further to become adults and were thus not able to reproduce further. Fig 3 shows a comparison between treated and untreated aphids. There was a clear difference between untreated and treated aphids after day 4, and by day 9 the treated aphids were all dead, while the untreated aphids were still alive.

**Fig 3.**
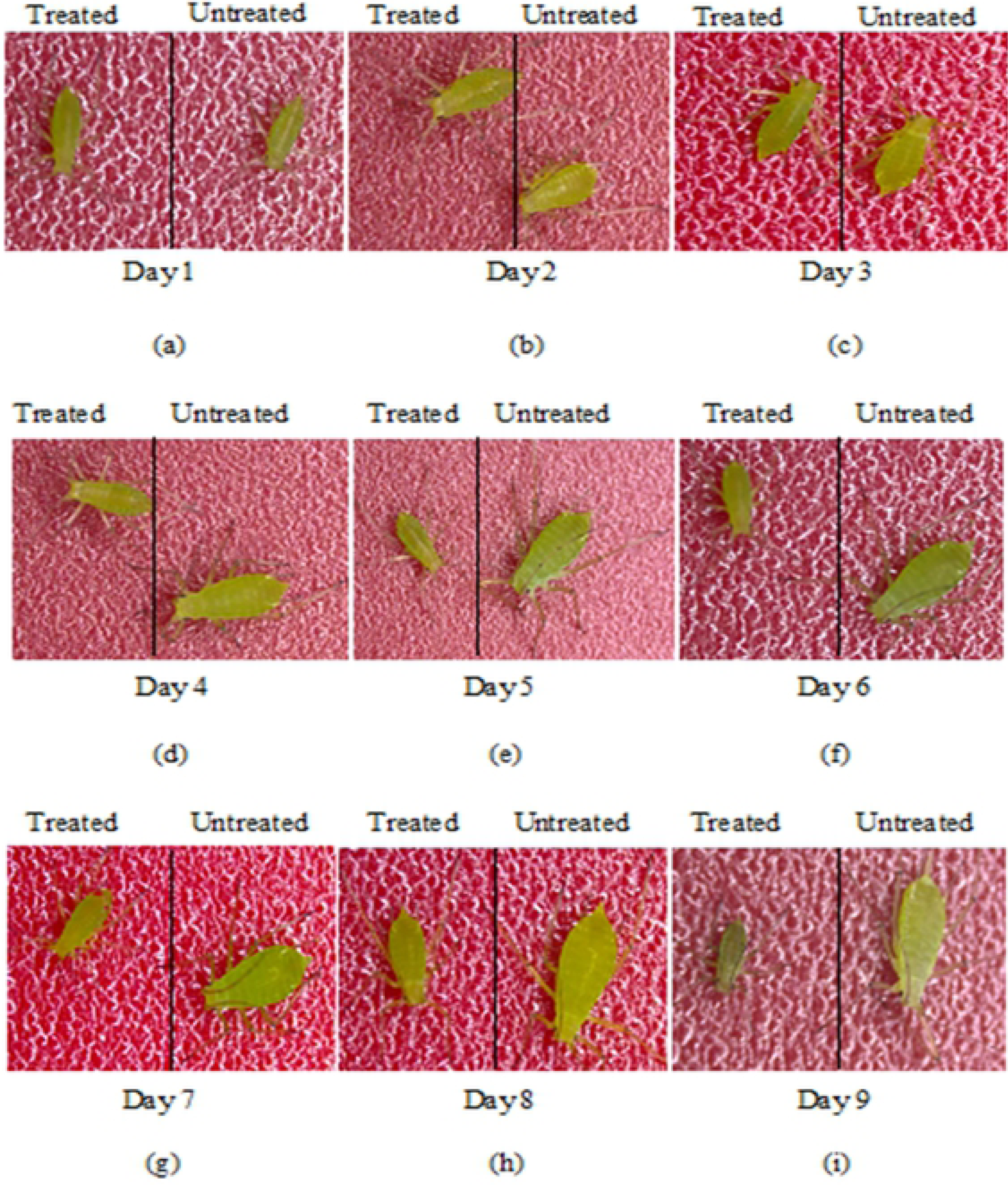
Comparison between growth of surviving aphids exposed to rosmarinic acid-treated and untreated diet after 24 h of bioassay. (a) to (i) comparison observed for up to 9 days, all treated aphids died by day 9

## Discussion

Screening candidate plants, purifying active ingredients, isolating and identifying the active plant constituents is required to discover new bioactive natural products [33]. We applied this methodology to identify rosmarinic acid as an active principle from the plant *I. rugosus.* Based on our previous study on the insecticidal activity of botanical extracts from various plant species, we found that the extract from *I. rugosus* was the most toxic to *A. pisum* [27] Further fractionation showed that the butanol fraction most likely contained the active principle. In this study we used the bioactivity-guided strategy to isolate and identify the active compound as rosmarinic acid. This strategy is interesting and has been used in previous studies to identify bioactive compounds. For example, the butanol fraction from *Citrullus colocynthis* was reported to be active against the black legume aphid, *Aphis craccivora*, and through the bioactivity-guided isolation strategy, the active principle, 2-O-ß-D-glucopyranosylcucurbitacin E, was successfully isolated [34]. Similarly, in another study involving bioactivity-guided isolation, the active principle, ailanthone, was isolated from the aqueous fraction of *Ailanthus altissima* against *A. pisum* [35].

In this study, the butanol fraction was subfractionated through reversed-phase flash chromatography. After bioactivity testing of all the resulting subfractions (1A-14A) against *A. pisum*, fraction 3A with lower LC values was selected for further fractionation. Through prep-LC, fraction 3A was subfractionated and the resulting subfractions (3A-1, 3A-2 and 3A-3) were analyzed for their bioactivity. Fraction 3A-3 with lower LC values was subjected to spectroscopic analysis. 1H NMR spectroscopy confirmed that the isolated fraction contained rosmarinic acid. However, due to the inconsistent chromatographic behavior during prep-LC, not enough compound could be collected to record 13C NMR data. The inconsistent chromatographic behavior with peak splitting observed could have arisen from several causes; a contamination on guard or analytical column inlet, a blocked frit or a small void at the column inlet (∼wear). The problem of peak shifting (variable retention times) could have been due to small changes in mobile composition, temperature fluctuations, column overloading or a combination of these problems which could have led to different UV patterns for each run. Due to this problem, the reversed-phase flash chromatography was repeated with a larger amount of the butanol fraction. Out of all the resulting subfractions (1B-6B), 1B was selected with lower LC values against *A. pisum*. Fraction 1B was subjected to acidic extraction to get two phases, aqueous and ethyl acetate. The ethyl acetate phase fraction was more active with lower LC values. After removing ethyl acetate, the active principle was identified through different spectroscopic techniques as rosmarinic acid. Similarly in another study, Chakraborty et al, [36] reported the isolation of caffeic acid and rosmarinic acid from *Basilicum polystachyon* through acidic extraction with HCl followed by partitioning with ethyl acetate and analyzed their antimicrobial activities.

This study reports the isolation and purification of rosmarinic acid (RA) from *I. rugosus* and its bioactivity against *A. pisum* for the first time. There was no significant difference observed between the bioactivity depicted by both isolated and commercial rosmarinic acid. *I. rugosus* rosmarinic acid gave LC values of LC_50_ = 0.2 ppm and LC_90_ = 5.4 ppm. These are very low LC values depicted after 24 h of bioassay and such low LC values have not been previously reported in any studies with compounds against *A. pisum* using the same feeding bioassay methodology [30, 37-39, 28, 40, 41]. This means that a very low amount of rosmarinic acid can cause significant toxic effects against *A. pisum* in 24 h. Very few insecticidal activities have been reported for rosmarinic acid. Regnault-Roger et al, [42] investigated the insecticidal activities of polyphenolic compounds, isolated from five plants belonging to Lamiaceae family against *Acanthoscelides obtectus* (Say) and observed that among all the polyphenolic compounds, rosmarinic acid and luteolin-7-glucoside were more toxic. An interesting avenue to follow for future studies will be the analyses of the underlying molecular mechanisms responsible for the cause of mortality in rosmarinic acid-treated aphids.

Additionally, a comparison between the growth of surviving aphids exposed to rosmarinic acid-treated and untreated diet after 24 h of bioassay was analyzed. It was clearly observed that the growth of surviving *A. pisum* nymphs stopped after 48 h of exposure to rosmarinic acid-treated diet, resulting in a size reduction and ultimately death as compared to aphids exposed to an untreated diet. A similar observation was made by Sadeghi et al, [30] who observed that the aphid size was reduced after 48 h of exposure to novel biorational insecticides, flonicamid and pymetrozine, and mortality was observed after 72 h.

## Conclusion

In this study, *I. rugosus* was identified as an interesting source for a botanical insecticide against *A. pisum*. Following bioactivity-guided selection, rosmarinic acid was isolated and identified through spectroscopic analysis as the bioactive compound in the *I. rugosus* extract. Based on the bioassay results, either the extracts from *I. rugosus* or the isolated insecticidal compound, rosmarinic acid could be exploited to develop potent aphicides, because of the high mortality of aphids caused at very low rosmarinic acid concentrations. This potential botanical insecticide may fit well in integrated pest management programs designed to control aphids. Considering that *I. rugosus* is already used for medicinal purposes, it is expected to be safer compared to the current conventional pesticides used to control aphids. Also, rosmarinic acid is known to reduce genotoxic effects induced by chemicals, which is contrary to some currently used toxic synthetic pesticides that could induce genotoxic effects in consumers. While this study highlights the potential of *I. rugosus* as a possible biopesticide source against a notorious insect pest such as *A. pisum*, it also provides the basis for further exploration and development of a formulation for effective field application.

## Acknowledgments

The authors are highly thankful to the PhD scholar, Kleber Pereira at the Faculty of Bioscience Engineering, Ghent University, Ghent, for maintaining and taking care of aphid colonies in the laboratory. We are grateful to Dr. Irum Shehzadi and other colleagues at the Department of Environmental Sciences, COMSATS University Islamabad, Abbottabad Campus, Pakistan, for their helpful support throughout this research project.

## Author Contributions

SK, CT, EB, RA, NF, SM, GS and MS conceived and designed the research. SK conducted the experiments. GS and SM contributed new reagents and/or analytical tools. SK, CT and EB, MA analyzed the data. SK, CT, EB, SM, GS and MS wrote the manuscript. All authors read and approved the manuscript.

